# T-cell development and activation in humanized mice lacking mouse major histocompatibility complexes

**DOI:** 10.1101/2024.03.26.586814

**Authors:** Milita Darguzyte, Philipp Antczak, Daniel Bachurski, Patrick Hoelker, Nima Abedpour, Rahil Gholamipoorfard, Hans A. Schlößer, Kerstin Wennhold, Martin Thelen, Maria Garcia-Marquez, Johannes König, Andreas Schneider, Tobias Braun, Frank Klawonn, Michael Damrat, Masudur Rahman, Jan-Malte Kleid, Sebastian J. Theobald, Eugen Bauer, Constantin von Kaisenberg, Steven Talbot, Leonard Shultz, Brian Soper, Renata Stripecke

**Affiliations:** Institute for Translational Immune-Oncology, Cancer Research Center Cologne-Essen (CCCE), University of Cologne, Cologne, Germany; University of Cologne, Faculty of Medicine and University Hospital Cologne, Department I of Internal Medicine, Center for Integrated Oncology Aachen Bonn Cologne Düsseldorf; Center for Molecular Medicine Cologne (CMMC), Cologne, Germany; Department II of Internal Medicine and Center for Molecular Medicine Cologne (CMMC), University of Cologne, Faculty of Medicine and University Hospital Cologne, Cologne, Germany; Cologne Cluster of Excellence on Cellular Stress Responses in Ageing-Associated Diseases, Cologne, Germany; Mildred Scheel School of Oncology Aachen Bonn Cologne Düsseldorf, Faculty of Medicine and University Hospital of Cologne, Cologne, Germany; Department of Translational Genomics, Faculty of Medicine and University Hospital Cologne, University of Cologne, Cologne, Germany; Center for Molecular Medicine Cologne (CMMC), University of Cologne, Faculty of Medicine and University Hospital Cologne, Cologne, Germany; Department of General, Visceral, Cancer and Transplantation Surgery, University of Cologne, Faculty of Medicine and University Hospital Cologne, Cologne, Germany; Department of Hematology, Oncology, Hemostasis and Stem Cell Transplantation, Hannover Medical School, Hannover, Germany; Department of Computer Science, Ostfalia University of Applied Sciences, Wolfenbuettel, Germany; Biostatistics Group, Helmholtz Centre for Infection Research, Braunschweig, Germany; Division of Infectious Diseases, Department I of Internal Medicine, University Hospital of Cologne, Cologne, Germany; German Center for Infection Research (DZIF), Partner Site Bonn-Cologne, 50931 Cologne, Germany; Institute of Transfusion Medicine, University of Cologne, University of Cologne, Faculty of Medicine and University Hospital Cologne, Cologne, Germany; Department of Obstetrics, Gynecology and Reproductive Medicine, Hannover Medical School, Hannover, Germany; Institute for Laboratory Animal Science, Hannover Medical School, Hannover, Germany; The Jackson Laboratory, Bar Harbor, ME, U.S.A.

**Keywords:** NSG, humanized mice, MHC, HLA, T-cell, IFN, lentivirus, immunization

## Abstract

Humanized mice transplanted with CD34^+^ hematopoietic progenitor cells (HPCs) are used to study human immune responses *in vivo*. However, the mismatch between the mouse major histocompatibility complexes (MHCs) and the human leukocyte antigens (HLAs) is not optimal for T-cell development and can trigger xenograft reactivity. We evaluated human T-cell development in NOD.Scid.Gamma mice lacking expression of MHC class I and II (NSG-DKO). Human leukocyte engraftment was detectable at 8 weeks post-transplantation. Human CD4^+^ and CD8^+^ T-cells were detectable in blood, thymus, bone marrow and spleen of humanized NSG-DKO mice for up to 20 weeks post-transplantation. Further, we evaluated the effects of lentiviral vector (LV) systemic delivery of HLA-A*02:01, HLA-DRB1*04:01, human GM-CSF/IFN-α and the human cytomegalovirus gB antigen. LV delivery promoted development and activation of human central memory, αβ and γδ T-cells with amplifications of the T-cell repertoire. LV administration unleashed multiple reactome pathways such as type-I interferon responses, cell cycle and metabolic processes. In summary, development of human T-cells in humanized mice does not rely on mouse MHCs and can be boosted systemically via LV administration.

## Introduction

Major Histocompatibility Complexes (MHCs) were discovered over a century ago in mice through the elucidation of the roles of genetic determinants of the immune system in both transplantation and cancer development (for a review see Götze & Burger, 1986). The discovery of the more genetically diversified human MHC homolog, the Human Leukocyte Antigen (HLA), enabled serological assays for optimizing the matching between donors and recipients of solid organ transplantation (for a review see Thorsby, 2009). In follow-up, hematopoietic stem cell transplantation (HCT) with bone marrow from HLA-matched donors into patient recipients with leukemia after high-dose radiation/chemotherapy resulted in engraftment, chimerism and eventual graft-versus-leukemia effects (for a review see Granot & Storb, 2020). As a result, HCT is nowadays an established curative clinical procedure for patients with hematologic malignancies or hematologic genetic defects.

Intriguingly, transplantation of irradiated immune deficient mice with enriched human hematopoietic stem cells (HSCs) or human progenitor cells (HPCs) result in mouse/human chimeric hematopoiesis and a semi-functional human immune system (HIS) (Anjos-Afonso & Bonnet, 2023). NSG mice combine the nonobese diabetic (NOD) inbred genetic background with the *Prkdc^scid^*mutation causing severe combined immunodeficiency and harbor a complete null mutation of the common cytokine receptor gamma chain (*Il2rγ*^null^). Transplantation of these mice with purified cord blood (CB) derived CD34^+^CD38^-^ HSCs achieves high and prolonged human chimerism with an effective development of T-cells displaying MHC-restricted cytotoxic functions (Ishikawa *et al*, 2005). Further, immune deficient radio-resistant NRG mice (combining the NOD and *IL2rγ^null^* stocks and incorporating targeted mutations in the recombination activating gene-1, *Rag1^null^*) showed comparable humanization with CD34^+^ HPCs, but less irradiation-induced tissue damage and xeno-graft-versus-host disease (xeno-GvHD) than NSG mice (Pearson *et al*, 2008).

Currently, several immune deficient mouse strains transplanted with CD34^+^ HPCs provide relevant models for elucidation of the human immune system, infectious disease and cancer immune-biology, and are becoming promising platforms for testing human-specific immune therapies (Saito *et al*, 2020; Chuprin *et al*, 2023). Although humanized mice recapitulate several aspects of HCT in humans, there is a scarce and delayed development of human T-cells. The lack or low levels of human cytokines, poor innate immune cell development and the underdeveloped lymphatic structures are some of the issues identified contributing to a weak development of human T-cells (Allen *et al*, 2019). Some of these problems were ameliorated when we applied human induced dendritic cells (iDCs) expressing human GM-CSF/IFN-α and antigens to humanized NRG (huNRG) mice (Salguero *et al*, 2014; Daenthanasanmak *et al*, 2015). Administration of iDCs promoted lymphatic regeneration and antigen-specific functional T- and B-cell immune responses against human cytomegalovirus (HCMV) pp65 and gB antigens (Volk *et al*, 2017; Theobald *et al*, 2020). However, we could not explain how mouse thymic tissues could promote human T-cell positive and negative selections resulting in polyclonal T-cells with functional T-cell receptors (TCRs).

Other groups demonstrated that humanization of transgenic NSG- and NRG-derived strains expressing human class I and class II MHCs/HLAs, as expected, showed improvements in T-cell development and cytotoxic functions (Najima *et al*, 2016; Danner *et al*, 2011; Majji *et al*, 2016; Masse-Ranson *et al*, 2019; Suzuki *et al*, 2012). Due to the difficulties of generating transgenic mice with several different HLA alleles, we explored lentiviral vector (LV) *in vivo* delivery of HLA-DRB1*04:01 (DR4) into huNRG mice (Kumar *et al*, 2021). Upon co-administration with LV expressing human GM-CSF/IFN-α and gB, we observed activation of T-cell and B-cell development (Kumar *et al*, 2021).

Here, we evaluated human T-cell development in humanized NSG-DKO (huNSG-DKO) mice lacking expression of class I and II mouse MHCs (Brehm *et al*, 2019). Further, we tested if LV-mediated gene delivery of HLA-DR4, HLA-A*02:01 (A2.1), human GM-CSF/IFN-α and gB would improve human T-cell development in the NSG-DKO background. We explored combinatory analyses of the human reconstitution using multicolor flow cytometry, high dimensional mass cytometry and mRNA sequencing analyses. Our data demonstrate that expression of MHCs in the mouse host tissues is dispensable for T-cell development. In addition, huNSG-DKO mice administered with LVs introducing human HLAs, human cytokines and a viral antigen showed a broad development of human central memory, αβ and γδ T-cells and activation of several immune and antiviral pathways. Therefore, *in vivo* LV delivery improves human adaptive immunity in our novel huNSG-DKO mouse model.

## Results

### Sub-lethally irradiated NSG-DKO can be humanized and show faster and higher human immune reconstitution in blood and bone marrow in comparison with huNRG mice

We performed triplicate humanization experiments using CD34^+^ cells isolated from three different CB donors (**Figure 1A**). HPCs from three different donors were used (all were positive for HLA-A*02:01 but none positive for HLA-DRB1*04:01, **Suppl. Table 1**) for transplantation of 5-7 weeks old mice after sub-lethal irradiation. The first goal was to evaluate if NSG-DKO mice irradiated with 150 cGy could be engrafted with human HPCs and could maintain human immune reconstitutions long-term. Radiation-resistant NRG mice irradiated with 450 cGy were used as our humanization reference control strain. Animals were routinely monitored for well-being according to score sheets with defined human endpoints that were approved by the animal welfare institutional office . 5-7-week-old NSG-DKO mice were smaller than NRG mice, but gained weight during the experiments (**Figure EV1A)**. Circulating human cells in blood were measured by flow cytometry on weeks 8, 12, 16 and 20 after HCT (antibodies used in the study are shown in **Suppl. Table 2**). Human lymphocytes homing in lymphatic tissues (bone marrow, thymus, spleen) were analyzed on week 20 after HCT (**Figure 1A; gating strategies are shown in Figure EV2A**). One additional cohort of humanized NSG-DKO mice was injected intravenously (i.v.) with LVs (the reconstitution analyses are described in more detail below). Longitudinal bioluminescence imaging (BLI) analyses of huNSG-DKO+LV showed biodistribution of bioluminescence signals (resulting from the expression of the luciferase transgenes) into anatomic regions of liver, spleen and bones, while control animals that did not receive LV did not show any bioluminescence (**Figure 1B, Figure EV1B**). The bioluminescence signal increased from week 8 to 12 post-HCT in huNSG-DKO+LV indicating persistent expression of the LV encoded transgenes (**Figure EV1C**). Already at week 8 post-HCT, huNSG-DKO mice had significantly higher percentages of huCD45^+^ cells in blood than huNRG mice (percentage of huCD45^+^ cells in blood at week 8: 50.6 ± 32.9 % in huNSG-DKO Vs. 14.0 ± 10.5 % in huNRG) (**Figure 1C, Suppl. Table 3, 4**). With exception of week 16, significantly higher humanization was observed until week 20 for NSG-DKO than NRG (**Figure 1C, Suppl. Table 3, 4**). Initially, high levels of B-cells were seen in both groups, while T-cells became more conspicuous at weeks 16 to 20 post-HCT (**Figure 1C, Suppl. Table 3, 4**). Compared with huNRG mice, huNSG-DKO mice showed significantly higher percentages of huCD45^+^ cells in bone marrow at 20 weeks after HCT (84.8 ± 6.2 % in huNSG-DKO Vs. 56.4 ± 24.9 % in huNRG) (**Figure 1D**, absolute counts can be seen in **Figure EV2D, Suppl. Table 5, 6**). Nonetheless, the percentages of T-cells and CD34^+^ cells in bone marrow were comparable for both strains (**Figure 1D, gating strategies are shown in Figures EV2B, C**). These results demonstrated consistent high and long-term human hematopoietic and immune reconstitution in NSG-DKO mice.

**Figure 1.**
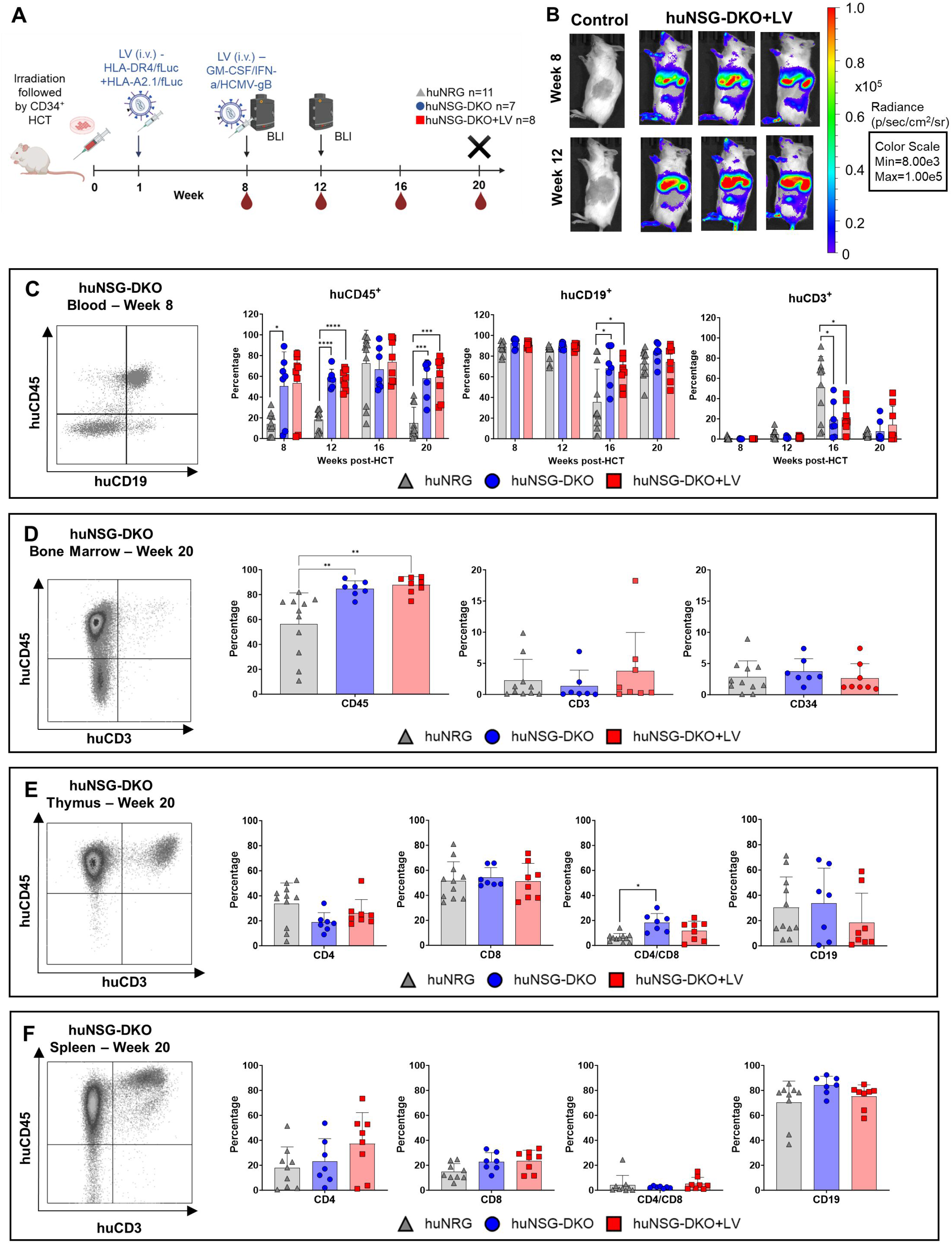
Human hematopoietic engraftment and T-cell development in humanized NSG-DKO mice. A Scheme of experiments: CD34^+^ stem cell transplantation (HCT) i.v. after irradiation, lentivirus (LV) immunization i.v., bioluminescence imaging (BLI) analyses, blood collections (BL) and termination (X). Cohorts are humanized huNRG, huNSG-DKO, huNSG-DKO+LV. The humanized mice were humanized with three different CB donors, both female and male mice were used in this experiment (details in **Suppl. Table 1**). B Full-body BLI quantified as photons/second (p/s) at 8 or 12 weeks post-HCT of huNSG-DKO control (representative one mouse) or after LV administration (representative three mice). C Quadrant on the left: FACS example showing development of B-cells. Graphs on the right: Blood analyses at weeks 8, 12, 16 and 20 after HCT and longitudinal quantification of cells expressing huCD45, huCD19 and huCD3 (in percentages). D Quadrant on the left: FACS example showing development of T-cells. Graphs on the right: Bone marrow analyses showing quantification of cells expressing huCD45, huCD3 or huCD34 (in percentages). E Quadrant on the left: FACS example showing development of T-cells. Graphs on the right: Thymus analyses and quantification of cells expressing huCD4, huCD8, huCD4/CD8, huCD19 (in percentages). F Quadrant on the left: FACS example showing development of T-cells. Graphs on the right: Spleen analyses and quantification of cells expressing huCD4, huCD8, huCD4/CD8, huCD19 (in percentages). Data information: In this figure, for blood analysis we used at week 8 and 12: 11 huNRG (grey triangles), 7 huNSG-DKO (blue circles) and 8 huNSG-DKO+LV (red squares). For analysis at week 16 and 20: 3 huNRG, 4 huNSG-DKO and 4 huNSG-DKO+LV. For thymus and bone marrow analysis we used: 11 huNRG, 7 huNSG-DKO and 7 huNSG-DKO+LV. For spleen analysis we used: 9 huNRG, 7 huNSG-DKO, 8 huNSG-DKO-LV. Statistical significances were determined by Wilcoxon-Mann-Whitney test with Bonferroni-Holm correction. Results are shown as mean +/-SD.

### Human T-cell development is not impaired in huNSG-DKO mice

Given the relevance of the MHC for T cell education and development in thymus, we assessed if CD3, one of the molecules of the TCR complex, was detectable in huNSG-DKO. Human CD3^+^ T-cells were detectable in thymus of huNSG-DKO mice (**Figure 1E**). In detail, analyses of thymus showed slightly lower percentages (but not lower total counts) of CD3^+^CD4^+^ T-cell in huNSG-DKO mice than in huNRG (**Figure 1E**, absolute counts can be seen in **Figure EV2E**). The percentages and total counts of CD8^+^ T-cells were similar between the humanized mouse strains. The double positive (huCD4^+^/huCD8^+^) T-cells were significantly magnified in huNSG-DKO compared with huNRG mice (percentage of huCD4^+^/huCD8^+^ cells in thymus: 18.3 ± 7.3 % in huNSG-DKO Vs. 6.2 ± 3.3 % in huNRG, **Figure 1E**, **Suppl. Table 5, 6**). Notably, for both humanized strains, CD19^+^ B-cells corresponded to about one third of the cells in the thymus (percentage of huCD19^+^ cells in thymus: 33.5 ± 28.0 in huNSG-DKO Vs. 30.3 ± 24.2 in huNRG, **Figure 1E**, **Suppl. Table 5, 6**). On the other hand, the analyses of CD3^+^ T-cells in spleens showed overall comparable frequencies of huCD4^+^, huCD8^+^ and double positive T-cells for huNRG and huNSG-DKO mice (**Figure 1F**, absolute counts can be seen in **Figure EV2F**), but CD19^+^ B-cells were largely over-represented in spleen (percentage of huCD19^+^ cells: 84.0 ± 7.3 in huNSG-DKO Vs. 70.2 ± 17.2 in huNRG, **Figure 1F, Suppl. Table 5, 6**) . Therefore, T-cells were observed at generally comparable levels in thymus and spleen of huNRG and huNSG-DKO, indicating that the mouse MHCs were dispensable for human T-cell education. Noteworthy, human B-cells, known to be relevant antigen presenting cells of the immune system, were detectable in high frequencies in both thymus and spleen of the humanized mice.

### Mainly memory phenotype T-cells are found in huNSG-DKO mice

In order to further assess T-cell maturation at 20 weeks post HCT, naïve, central memory, effector memory and terminal effector phenotypes were enumerated by flow cytometry (see detailed gating strategy in **Figure EV3A**). In all analyzed tissues (bone marrow, thymus and spleen) mainly effector memory T-cells were found (central memory and effector memory T-cells can be seen in **Figure 2A**, **C, E**, naïve and terminal effector in **Figure EV3B, C, D**). The frequency of effector memory T-cells was in general higher in huNSG-DKO compared to huNRG (percentage of effector memory huCD4^+^ T-cells in spleens: 48.2 ± 18.5 in huNSG-DKO Vs. 29.1 ± 19.3 in huNRG, **Figure 2E**, **Suppl. Table 5, 6**). Moreover, activation in regard to the expression levels of PD-1 and CD69 within huCD4^+^ and huCD8^+^ T-cells was analyzed (see detailed gating strategy in **Figure EV3A**). T-cells of huNSG-DKO mice showed in general higher PD-1 expression (**Figure 2B, D, F**). For example, in thymus, PD-1 expression in CD8^+^ T-cells of huNSG-DKO was three-fold higher compared to huNRG mice (MFI of PD-1 on huCD8^+^ in thymus: 51,405 in huNSG-DKO vs 16,244 huNRG, **Figure 2D**, **Suppl. Table 5, 6**) . Expression of CD69, reflecting activation, was in general higher for T-cells of huNSG-DKO mice (**Figure 2B, D, F**). Analyses of B-cell development and maturation markers (see detailed gating strategy in **Figure EV3E**), showed significantly higher absolute counts of naïve cells and plasmablasts in huNSG-DKO compared to huNRG mice (IgD^+^CD27^-^ B-cell counts in spleen: 8,024,367 in huNSG-DKO vs. 3,443,476 in huNRG, **Figure 2G**, **Suppl. Table 5, 6**), but the levels of IgM^+^, IgG^+^ and IgA^+^ cells were comparable between the strains (**Figure EV3F**). Thus, as a trend, activation of T- and B-cells was moderately higher in lymphocytes of huNSG-DKO mice, indicating functional immune reactivities.

**Figure 2.**
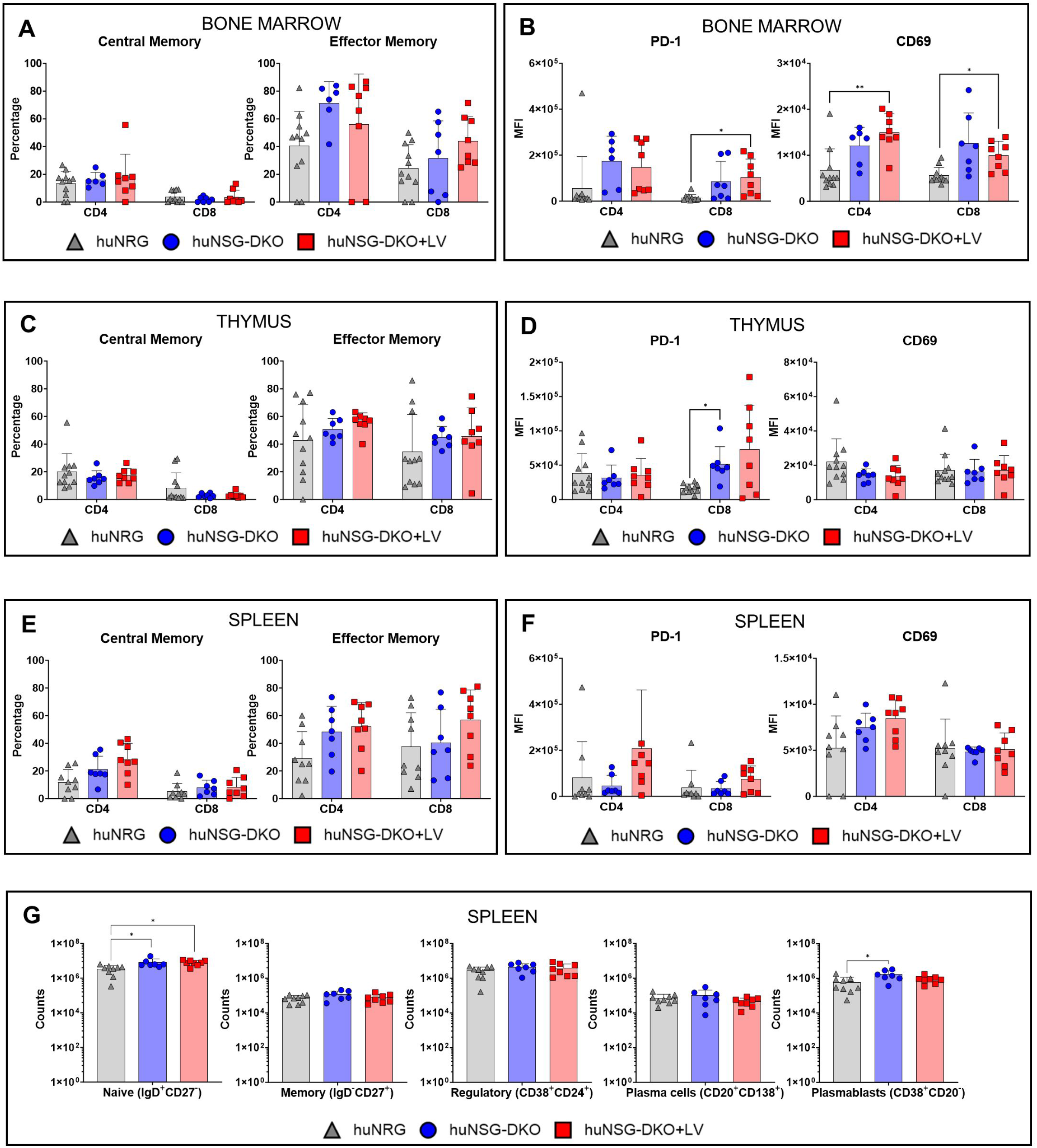
Human T-cell and B-cell maturation and activation in humanized NSG-DKO mice. A Analysis of Central Memory and Effector Memory T-cell subtypes within huCD4^+^ and huCD8^+^ T-cells in bone marrow (in percentages). B Analysis of T-cell activation markers PD-1 and CD69 within huCD4^+^ and huCD8^+^ T-cells in bone marrow (in absolute numbers, log scale). C Analysis of Central Memory and Effector Memory T-cell subtypes within huCD4^+^ and huCD8^+^ T-cells in thymus (in percentages). D Analysis of T-cell activation markers PD-1 and CD69 within huCD4^+^ and huCD8^+^ T-cells in thymus (in absolute numbers, log scale). E Analysis of Central Memory and Effector Memory T-cell subtypes within huCD4^+^ and huCD8^+^ T-cells in spleen (in percentages). F Analysis of T-cell activation markers PD-1 and CD69 within huCD4^+^ and huCD8^+^ T-cells in spleen (in absolute numbers, log scale). G Analysis of B-cell subtypes in spleens. B-cell subtypes: naïve, memory, regulatory, plasma cells, plasmablasts, (in absolute numbers, log scale). Data information: In this figure, for analysis of thymus and bone marrow we used: 11 huNRG, 7 huNSG-DKO and 7 huNSG-DKO+LV. For spleens we used: 9 huNRG (grey triangles), 7 huNSG-DKO (blue circles), 8 huNSG-DKO+LV (red circles). For spleen analysis we used: 9 huNRG, 7 huNSG-DKO, 8 huNSG-DKO-LV. Statistical significances were determined by Wilcoxon-Mann-Whitney test with Bonferroni-Holm correction. Results are shown as mean +/-SD.

### *In vivo* LV delivery promotes T-cell development in huNSG-DKO

Within our experimental design, we wanted to evaluate if immune responses in huNSG-DKO could be enhanced. As an additional cohort for the huNSG-DKO mice, we evaluate in parallel the effects of *in vivo* delivery of multicistronic LVs to deliver HLAs (LV-HLA-DR4/fLuc and LV-HLA-A2.1/fLuc) and to stimulate immune responses (LV-hGM-CSF/ hIFN-α/ HCMV-gB). LVs expressing HLAs were administered i.v. one-week post-HCT, while the immune stimulatory LV was given eight weeks post-HCT (the schemes of the LV multicistronic constructs are shown in **Figure EV4A,** expression of DR4 and A2.1 in transduced 3T3 mouse fibroblasts is shown in **Figure EV3B**). No signs of distress were observed after LV administration, as body weight increased during the course of experiments indicating well-being of animals and no signs of xeno-GvHD (**Figure EV1A**). Expression of fLuc was confirmed by BLI analyses in anatomic regions of liver and spleen of huNSG-DKO (**Figure 1B**) and non-humanized NSG-DKO mice (**Figure EV1B, C**). HuNSG-DKO mice administered with LVs showed an enhancement in the percentages of T-cells in blood at week 20 (percentage of huCD3^+^ cells : 14.2 ± 18.4 in huNSG-DKO+LV Vs. 7.5 ± 10.7 % in huNSG-DKO, **Figure 1C, Suppl. Table 3, 4**). Analyses of bone marrow (**Figure 1D**), thymus (**Figure 1E**) and spleen (**Figure 1F**) also indicated a perceptible enhancement of the CD4^+^ and/or CD8^+^ T-cell development after LV administrations compared with huNSG-DKO mice. The effects of LV administration on T-cell phenotype and activation were further investigated. In spleen, accumulation of terminal effector T-cells were seen upon *in vivo* LV delivery (percentage of effector memory huCD4^+^ T-cells in spleen: 48.2 ± 18.5 % in huNSG-DKO, and 52.2 ± 17.3 % in huNSG-DKO+LV, **Figure 2E**, **Suppl. Table 5, 6**). The same trend was observed regarding higher T-cell activation of huCD8^+^ T-cells in thymus in response to LV administration (MFI of PD-1: 51,405 in huNSG-DKO and 73,208 in huNSG-DKO+LV, **Figure 2C**, **Suppl. Table 5, 6**). Regarding CD4^+^ T-cell activation analyses via CD69 upregulation, modest effects were observed in bone marrow (MFI of CD69: 12,055 in huNSG-DKO and 14,995 in huNSG-DKO+LV, **Figure 2A**, **Suppl. Table 5,6**). Analyses of splenic B-cells showed similar immunophenotypic patterns of cells obtained from huNSG-DKO or huNSG-DKO+LV mice (**Figure 2G, Figure EV3F**). Thus, an immune modulation of T-cells after LV administration was seen in huNSG-DKO mice.

### Human NK, γδ T-cells, monocytes and dendritic cells identified by high dimensional cell clustering data

Fluorescence-based flow cytometry is the most widely adopted method to quantify the percentages and absolute counts of human immune cells in humanized mice. Nonetheless, the numbers of markers analyzed are limited to up to twenty parameters in conventional devices and sometimes the fluorochromes are difficult to compensate. A more advanced method allowing higher number of markers is the cytometry by time-of-flight (CyTOF) or mass cytometry that utilizes antibodies labeled with heavy metal isotopes. The resulting immune cell abundances are detected using a time-of-flight mass spectrometer. We developed a method to analyze untouched cryopreserved/thawed bone marrow samples (**Figure 3A**). As reference material to validate the method, we used human cryopreserved CB as a positive control and bone marrow recovered from non-humanized NRG and NSG-DKO mice as negative controls (**Figure EV5A**). For these analyses, we included mice from a single CB donor humanization. For each mouse 1×10^6^ total cells were used for labeling (antibodies used for labeling are shown in **Suppl. Table 7**). By employing a human CD45^+^ barcoding step, the mouse cells were excluded from further analysis steps. We checked 30 markers simultaneously per sample and the bioinformatics analyses were performed for high-dimensional analyses by marker expression distribution across all events. Dimensionality reduction used t-distributed stochastic neighbor embedding (t-SNE) and a clustering algorithm (FlowSOM) for visualization to identify and quantify the cellular heterogeneity among the three different experimental cohorts (**Figure 3A**). Labeling of human markers were not detectable in non-humanized NRG or NSG-DKO control mice (**Figure EV5A**), while ten major human cell compartments were identified in humanized mice (**Figure 3B, Figure EV5B**). Similar analyses by FACS (**Figure 1D**), CyTOF analyses showed that B-cells (early B precursor cells and mature B-cells) constituted the largest population in bone marrow (**Figure 3C**). Larger amounts of CD4^+^ T-cells were seen in huNSG-DKO+LV than in huNSG-DKO and huNRG mice (**Figure 3C, D**) (percentages of huCD4^+^ cells at week 20: 0.6 ± 0.5 % in huNRG, 2.8 ± 4.6 % in huNSG-DKO and 7.3 ± 6.4 % in huNSG-DKO+LV, **Figure 3D, Suppl. Table 8, 9**). Further, CyTOF identified human monocytes as an abundant population in all cohorts (**Figure 3C, D, Suppl. Table 8, 9**), but with increased percentages in huNSG-DKO+LV compared with huNSG-DKO. Interestingly, this was paired with higher frequencies of plasmacytoid dendritic cells (pDCs) and myeloid dendritic cells (mDCs) in huNSG-DKO+LV mice (**Figure 3B, C, D**). Further, natural killer (NK) and γδ T-cells were detectable in all cohorts but some of the huNSG-DKO mice that received LV showed outliers with much higher frequencies (percentage of γδ T-cells: 0.04 ± 0.02 % in huNRG, 0.2 ± 0.2 % in huNSG-DKO and 0.4 ± 0.4 % in huNSG-DKO+LV, **Figure 3C, D** **, Suppl. Table 8, 9**). These results indicated stimulatory effects of the LV administration in the innate immune cells of bone marrow. Moreover, the activation of T-cells was evaluated by CD69 expression (**Figure 3E**). Comparably to FACS analysis (**Figure 2B**), huNSG-DKO+LV mice had the highest levels of activated T-cells (CD69 expression on huCD4^+^ T-cells in bone marrow: 0.442 in huNRG, 0.550 in huNSG-DKO and 0.592 in huNSG-DKO+LV, **Figure 3E, Suppl. Table 8, 9**). HLA-DR expression was particularly enhanced on CD8^+^ T-cells of the huNSG-DKO+LV mouse cohort (HLA-DR expression on huCD8^+^ T-cells in bone marrow: 0.125 in huNRG, 0.180 in huNSG-DKO and 0.236 in huNSG-DKO+LV, **Figure 3E, Suppl. Table 8, 9**), suggesting the presence of CD8^+^ regulatory T-cells in humanized mice (Machicote *et al*, 2018).

**Figure 3.**
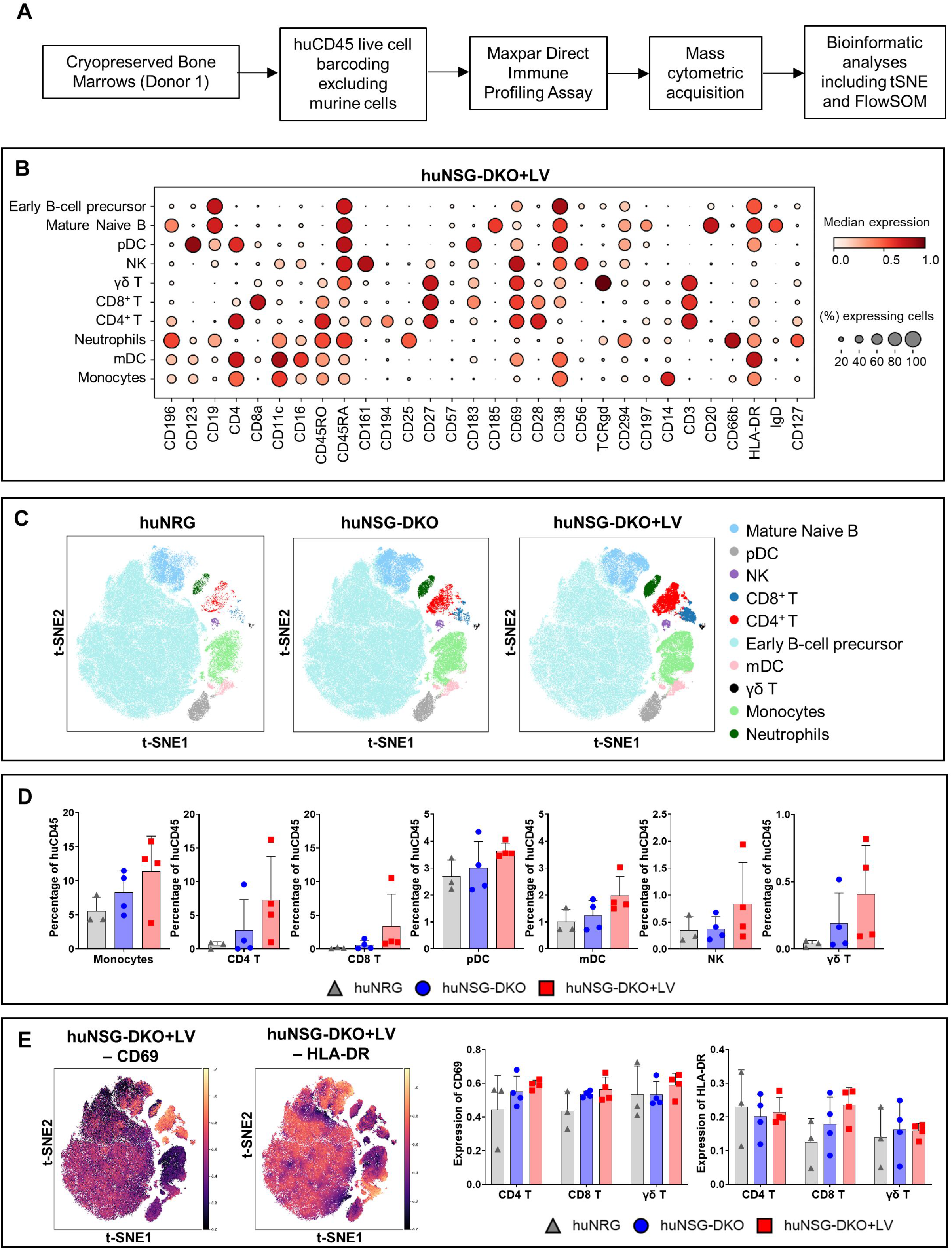
Humanized NSG-DKO mice develop NK cells and γδ T-cells. A Scheme of sample preparation, staining, CyTOF measurement and analysis of bone marrow samples. The mice were humanized using only one donor. B CyTOF analyses and dotplot of huNSG-DKO+LV cell subtypes (a total of 160,343 cells) and their expression of different Maxpar Direct Immune Profiling lineage markers. The dot size corresponds to the fraction of cells expressing the indicated marker within each cell type, and the color indicates median expression. C Anti-human CD45-CD live-cell barcoded analysis of immune cell types in bone marrow samples 20 weeks post-HCT. A total of 152,755 cells, 185,888 cells and 160,343 cells were analyzed for huNRG, huNSG-DKO, and huNSG-DKO+LV respectively. t-SNE plots displaying different subtypes of human immune cells clustered by FlowSOM and annotated manually using the lineage markers presented in the dot plot (panel B). D CyTOF analysis of CD4^+^, CD8^+^ and γδ T-cells, natural killer, monocytes, myeloid and plasmacytoid dendritic cell counts in bone marrow samples (in percentages). E Overlay of CD69 and HLA-DR expression on t-SNE embeddings of huNSG-DKO+LV across various cell types, as depicted in panel B. CD69 and HLA-DR expression in CD4^+^, CD8^+^ and γδ T-cells (in absolute numbers). Data information: In this figure, we used for analyses bone marrows from mice humanized using same donor: 3 huNRG, 3 huNSG-DKO, 4 huNSG-DKO+LV. Statistical significances were determined by Wilcoxon-Mann-Whitney test with Bonferroni-Holm correction. Results are shown as mean +/-SD.

### Identification of human biomarkers of response by mRNA sequencing analyses

To get a global idea about gene expression in humanized mice and the effects of LV delivery, we performed bulk mRNA sequencing of spleen cells (**Figure 4A**). Due to high dimensionality of the data we decided to restrict the samples to mice humanized with one CB donor. Since humanized mice possess genetic material from both mice and human, the assignment of each identifiable transcript had to be assessed to the two species. For this assignment, control samples of non-humanized mice possessing only mouse genes and huPBMCs possessing only human genes were also sequenced. Since mouse and human have a high genetic similarity, it was important to identify potential genes where the aligner would erroneously assign a read to the wrong species (Chinwalla *et al*, 2002). These genes were removed from further analysis. To visualize all of the data, a heatmap was generated of all 59,314 genes analyzed across the two species (**Figure 4B**). It shows that the majority of genes are specifically assigned to the right species. Control samples such as the non-humanized mouse samples show a generally higher expression of mouse genes as compared to the other models and an extremely low value for human genes as no expression of these genes is expected here. As expected, mouse genes were identified near or below the detection limit in huPBMCs and human genes expressed at higher levels (**Figure 4B**). Similarly, several human transcripts could be identified in mRNA obtained from humanized mouse samples compared with non-humanized mice. Close comparison of two humanized mouse strains revealed that huNSG-DKO mice have more human and less mouse genes upregulated than huNRG (**Figure 4C**), corresponding to the better humanization in the NSG-DKO strain.

**Figure 4.**
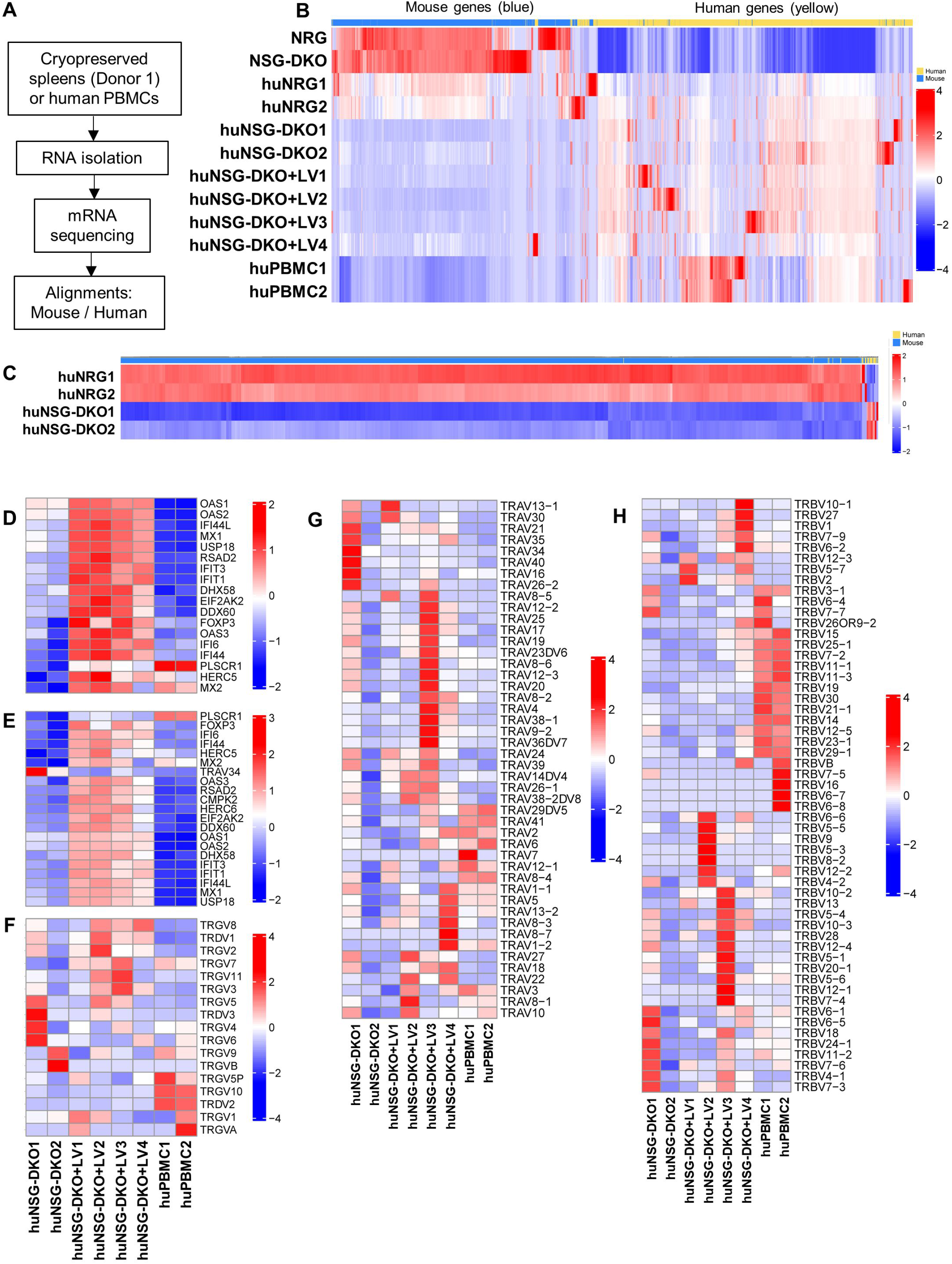
Humanized NSG-DKO mice show upregulation of several biomarkers of immune responses after LV delivery. A Scheme of sample preparation, RNA isolation, mRNA sequencing and analysis of spleen samples. The mice were humanized using only one donor. B Alignment of mouse (blue) versus human (yellow) genes. As expected, non-humanized NRG and NSG-DKO mice have higher frequencies of mouse upregulated transcripts than humanized mice and PBMCs. HuNRG mice still upregulate some mouse genes, while in comparison huNSG-DKO mice do not. C Differentially expressed genes between humanized NRG and NSG-DKO mice. Higher frequencies of mouse transcripts are seen in huNRG, while huNSG-DKO upregulate more human genes. D Genes responsible for defense response to viruses are upregulated in huNSG-DKO+LV. E Genes responsible for response to other organism are upregulated in huNSG-DKO+LV. F Recombinations in T-cell receptor G and D variable chains are polyclonal and more frequent in huNSG-DKO+LV. G Recombinations in T-cell receptor A variable chain are polyclonal and more frequent in huNSG-DKO+LV. H Recombinations in T-cell receptor B variable chain are polyclonal and more frequent in huNSG-DKO+LV. Data information: In this figure, we used for analyses spleens from mice humanized using same donor: 2 huNRG, 2 huNSG-DKO, 4 huNSG-DKO+LV. For controls 2 PBMCs and spleens of non-humanized (no HCT) mice 1 NRG and 1 NSG-DKO were used.

### Identification of human biomarkers and pathways of response to LV delivery

*In vivo* LV delivery introducing human HLAs, human cytokines and a viral antigen, showed some effects by flow cytometry and CyTOF on the human innate and adaptive compartments in huNSG-DKO. In order to confirm the effects of the LV delivery at the gene expression level, the genes involved in pathways identifying “defense against viruses” (**Figure 4D**) and “response to other organisms” (**Figure 4E**) were compared between huNSG-DKO with and without LV administration. It can be clearly seen that only mice injected with LV displayed high upregulation of several genes associated with immune responses, confirming our hypothesis. Among the top highest identifiable transcripts associated with interferon responses were *IFI44, IFIT3, IFIT1* and *IFI6* (**Table 1**). Moreover, transcripts encoding for proteins with additional immunological activities were *XAF1*, *CSF2RA* and *FOXP3* (**Table 1**). Several transcripts encoding for enzymes were identified, most of them interferon inducible and with antiviral activities (*MX1, USP18, OAS1, OAS2, OAS3*) (**Table 1**). Furthermore, the expression of TCR-γδ variable chains (**Figure 4F**), TCR-α variable chains (**Figure 4G**) and TCR-β variable chains (**Figure 4H**) were compared. In this case, although the differences between huNSG-DKO and huNSG-DKO+LV were less evident, mice administered with LV seem to exhibit larger TCR variable repertoires, i.e., higher polyclonality. A deeper look at the TCR transcripts that were significantly more abundant in huNSG-DKO+LV compared with huNSG-DKO highlighted one public clonotypes assigned to immune responses against Influenza (*TRBV19*), other public clonotypes (*TRBV13, TRBV1*) and alpha and beta joining sequences (**Table 1**). Besides looking at single transcriptomes, genes responsible for similar function were clustered and genome pathways or ‘reactomes’ were analyzed (**Figure 5**). Interestingly, multiple human genome pathways were significantly upregulated, particularly immune responses, response to type I interferon, type I interferon production, response to virus, negative regulation of viral genome replication, histamine biosynthetic process (**Figure 5**, **Table 2**). Interestingly, other pathways were also activated huNSG-DKO+LV in comparison to non-treated mice, such as actin filament network formation, regulation of ribonuclease activity and biological process involved in interspecies interaction between organisms (**Figure 5**, **Table 2**). Furthermore, we also interrogated which murine transcripts were more abundant in huNSG-DKO+LV in comparison to huNSG-DKO mice and we identified several genes associated with RNA and DNA metabolism (**Table 3**). Thus, several immune and non-immune pathways operate in huNSG-DKO that can be activated upon LV administration.

**Figure 5.**
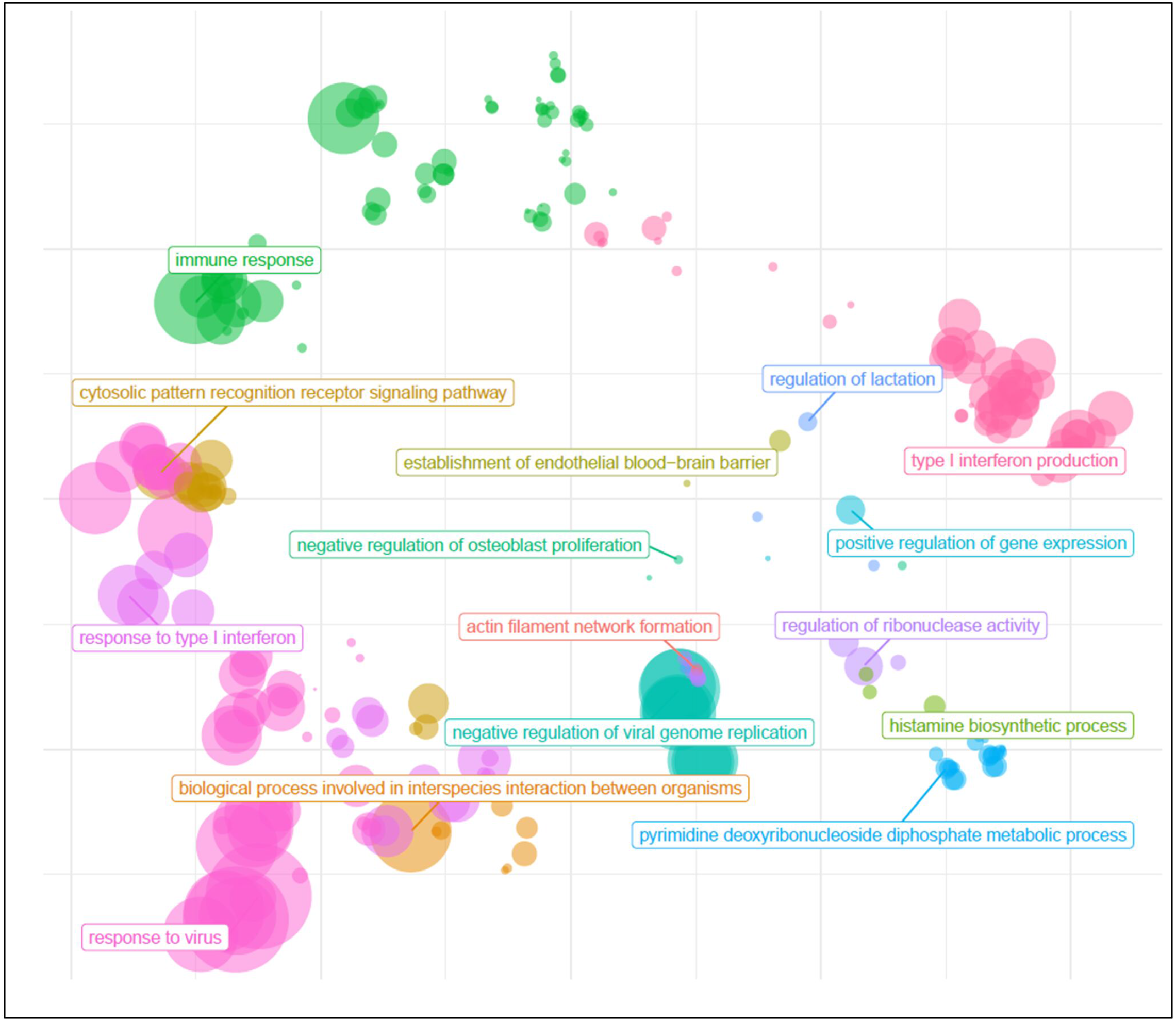
Humanized NSG-DKO mice show activation of several immune and non-immune-related pathways after LV delivery. The pathways are grouped and only parent terms that are based on the hierarchical nature of gene ontology are shown. Immunological pathways: Immune response, response to type I interferon, type I interferon production, response to virus, negative regulation of viral genome replication, histamine biosynthetic process. Other pathways: Actin filament network formation, regulation of ribonuclease activity, biological process involved in interspecies interaction between organisms, regulation of lactation, pyridine deoxyribonucleoside diphosphate metabolic process, negative regulation of osteoblast proliferation. The sizes of the bubbles represent their relevance.

**Table 1.** Human biomarkers associated with TCR sequences, immune responses and enzymes upregulated in huNSG-DKO+LV compared with huNSG-DKO.

| Biomarker Class | Gene Name | Description | Term ID | p value | Remarks |
| --- | --- | --- | --- | --- | --- |
| TCR Sequences | TRBV19 | T-cell receptor beta variable 19 | ENSG00000211746.3 | 2.02E-12 | Public clonotype, response to influenza (Yang *et al*, 2017) (Huisman *et al*, 2022) |
|  | TRBV10-2 | T-cell receptor beta variable 10-2 | ENSG00000229769.2 | 2.98E-09 | Predicted to be part of the TCR complex |
|  | TRBJ1-5 | T-cell receptor beta joining 1-5 | ENSG00000282173.1 | 2.99E-08 | J segment usage decreased in patients with Crohns disease (Hong *et al*, 2022) |
|  | TRAJ9 | T-cell receptor alpha joining 9 | ENSG00000211880.1 | 1.10E-07 | In germline T-cell receptor alpha and delta sequences |
|  | TRAJ48 | T-cell receptor alpha joining 48 | ENSG00000211841.1 | 4.66E-07 | In germline T-cell receptor alpha and delta sequences |
|  | TRBV1 | T-cell receptor beta variable 1 | ENSG00000242736.1 | 1.07E-06 | Pseudogene and public clonotype, decreased in patients with Crohns disease (Hong *et al*, 2022) |
|  | TRGV7 | T-cell receptor gamma variable 7 | ENSG00000249978.1 | 2.47E-05 | Pseudogene |
|  | TRBV13 | T-cell receptor beta variable 13 | ENSG00000276405.1 | 6.36E-05 | Public clonotype, predicted to be part of the TCR complex |
| Immune responses | IFI44L | Interferon induced protein 44 like | ENSG00000137959.17 | 9.85E-15 | Interferon induced, antiviral, macrophage response to bacterial infection |
|  | IFI44 | Interferon induced protein 44 | ENSG00000137965.11 | 1.11E-12 | Interferon induced, immune response |
|  | IFIT3 | Interferon induced protein with tetratricopeptide repeats 3 | ENSG00000119917.15 | 3.35E-08 | Interferon induced, antiviral |
|  | IFIT1 | Interferon induced protein with tetratricopeptide repeats 1 | ENSG00000185745.10 | 5.15E-08 | Interferon induced, antiviral |
|  | XAF1 | XIAP associated factor 1 | ENSG00000132530.17 | 1.88E-07 | Antiviral, may be a tumor suppressor |
|  | CSF2RA | Colony stimulating factor 2 receptor subunit alpha | ENSG00000198223.18 | 2.74E-07 | Receptor for granulocyte-macrophage colony-stimulating factor (GM-CSF) |
|  | IFI6 | Interferon alpha inducible protein 6 | ENSG00000126709.16 | 5.69E-07 | Interferon induced, antiviral |
|  | FOXP3 | Forkhead box P3 | ENSG00000049768.18 | 4.61E-05 | Regulatory T-cell development and function |
| **Enzymes** | CMPK2 | Cytidine/uridine monophosphate kinase 2 | ENSG00000134326.12 | 3.59E-08 | Mitochondrial, antiviral, interferon-dependent and non-dependent activities |
|  | MX1 | MX dynamin like GTPase 1 | ENSG00000157601.15 | 7.49E-08 | Interferon induced, antiviral |
|  | USP18 | Ubiquitin specific peptidase 18 | ENSG00000184979.11 | 2.64E-07 | Interferon induced, antiviral |
|  | OAS2 | 2'-5'-oligoadenylate synthetase 2 | ENSG00000111335.14 | 7.60E-06 | Interferon induced, antiviral |
|  | OAS3 | 2'-5'-oligoadenylate synthetase 3 | ENSG00000111331.14 | 9.99E-06 | Interferon induced, antiviral |
|  | OAS1 | 2'-5'-oligoadenylate synthetase 1 | ENSG00000089127.15 | 1.77E-05 | Interferon induced, antiviral |
|  | HERC5 | HECT and RLD domain containing E3 ubiquitin protein ligase 5 | ENSG00000138646.9 | 2.40E-05 | Macrophage response to bacterial infection |
|  | DHX58 | DEXH-box helicase 58 | ENSG00000108771.13 | 3.25E-05 | Negative regulation of immune response and interferon response |

**Table 2.** Human pathways upregulated in huNSG-DKO+LV compared with huNSG-DKO.

| Term ID | Name | Intersection size | p value |
| --- | --- | --- | --- |
| GO:0051607 | Defense response to virus | 17 | 3.24E-19 |
| GO:0009615 | Response to virus | 18 | 3.24E-19 |
| GO:0006955 | Immune response | 22 | 6.88E-12 |
| GO:0044419 | Biological process involved in interspecies interaction between organisms | 21 | 6.88E-12 |
| GO:0043207 | Response to external biotic stimulus | 20 | 6.89E-12 |
| GO:0045071 | Negative regulation of viral genome replication | 8 | 6.89E-12 |
| GO:0051707 | Response to other organism | 20 | 6.89E-12 |
| GO:0009607 | Response to biotic stimulus | 20 | 9.19E-12 |
| GO:0140546 | Defense response to symbiont | 17 | 6.73E-11 |
| GO:0045069 | Regulation of viral genome replication | 8 | 1.45E-10 |

**Table 3.**
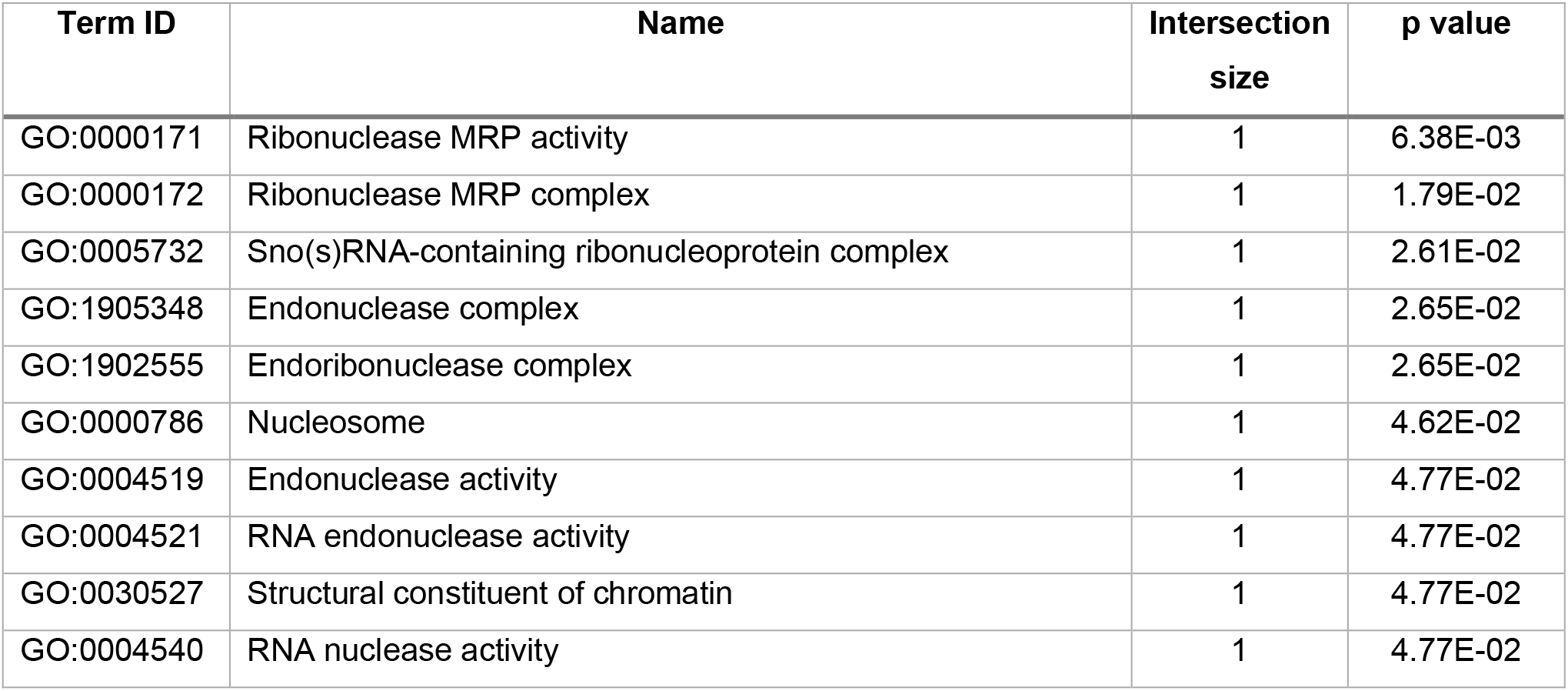
Murine pathways upregulated in huNSG-DKO+LV compared with huNSG-DKO.

## Discussion

Humanized mouse models have advanced in the last decades from basic science towards relevant preclinical models for translational research, particularly in the immunology field. As the demand for humanized mouse models grows to evaluate the potency and safety of human-specific immune-modulatory biologics (Waldron-Lynch *et al*, 2012; Yan *et al*, 2019a), several optimizations to enhance engraftment and development of human cells in HIS-mice are ongoing (Stripecke *et al*, 2020). One important novel contribution of NSG-DKO mouse strain humanized with PBMCs is the reduction of the xenoGvHD, associated with non-specific human T-cell reactivity against the mouse tissues. Counteracting xenoGvHD is key in order to conduct long-term studies to detect and mechanistically characterize *bona fide* immune toxicities, such as cytokine release syndrome (CRS) (Brehm *et al*, 2019; Ashizawa *et al*, 2017; Matas-Céspedes *et al*, 2020). Brehm et al. reported a more gradual and persistent human T-cell expansion in NSG-DKO mice compared to the parental NSG strain having both MHCs, but eventually xenograft reactivity occurred after several weeks (Matas-Céspedes *et al*, 2020).

On the other hand, fully humanized mouse models generated after CD34^+^ HPC transplantation can maintain functional human T-cells developed endogenously devoid of xeno-GvHD for more than 20 weeks (Theobald *et al*, 2020). Immunodeficient mice implanted with human fetal liver HSCs along with fetal thymus and infused with autologous fetal liver CD34^+^ cells (BLT mice) display good development of T-cells, B-cells and monocytes (Melkus *et al*, 2006). BLT mice have been broadly used for testing immunotoxicity via CRS (Yan *et al*, 2019b) and studying and developing drugs against HIV infection (Denton *et al*, 2012; Ling *et al*, 2023). However, due to limited availability of human fetal tissue and ethical concerns, other CD34^+^ sources such as mobilized peripheral blood or CB are presently more commonly used.

Currently, the basic knowledge regarding how implanted human HPCs differentiate and develop within these chimeric mice and result in MHC restricted human T-cells is still limited. It has been postulated that human T-cells are trained on mouse MHCs in thymus. Indeed knocking out murine MHC class I has produced less huCD8^+^ T-cells, while MHC class II knock out less huCD4^+^ T-cells, which seems to be in line with the proposed hypothesis (Pino *et al*, 2010; Läderach *et al*, 2023). In our study, NSG-DKO mice were humanized with CD34^+^ CB HPCs from three different donors. Although a limitation of our study is the modest cohort sizes, we obtained consistent humanization levels using CD34^+^ HPCs from three different CB donors. Interestingly, we did see human T-cell development in huNSG-DKO and, in fact, the overall humanization rate (amount of huCD45^+^ cells) was higher in huNSG-DKO in comparison to the huNRG mice. We acknowledge using NSG instead of NRG mice as control would have been more relevant reference strain. However, Pearson et al showed that NRG mice support similar levels of human lympho-hematopoietic cell engraftment as NSG mice (Pearson *et al*, 2008). Hence, we believe that the superior HIS reconstitution are mainly due to knock out of mouse MHCs in NSG-DKO mice. Although the explanation of this phenomenon is beyond the scope of this current work, one possibility is that due to the lack of mouse MHCs there are less inflammatory xenograft reactions already at the time of engraftment and the mouse myeloid cells are less activated to perform phagocytosis of the human cells.

We presumed that human T-cell development could have been impaired in NSG-DKO mice due to absence of MHC expression in the thymus epithelial cells (TECs). But it seems that other cells and lymphatic organs could take over T-cell education in NSG-DKO mice, just like it happens in humans who undergo thymectomy (Torfadottir *et al*, 2006; Cavalcanti *et al*, 2021). However, thymic T-cell development was not impaired and, in fact, our huNSG-DKO had even higher levels of thymic T-cells in comparison to huNRG mice. After having a closer look we identified that these were mainly effector memory T-cells. In the thymus mainly naïve T-cells are expected, but it has been observed that mature T-cells reenter the thymus (Naparstek *et al*, 1982). As it has been shown that these returning T-cells can contribute to positive and negative selection, we presume that this one of the ways how T-cells get selected in our mouse strain that lacks MHCs (King *et al*, 2009; Bosco *et al*, 2009). Additionally, it has been seen that memory CD8^+^ T-cells that reenter the thymus can protect the thymus against infections (Hofmann et al, 2013). We presume that huNSG-DKO mice with higher levels of huCD8^+^ T-cells in thymus would have a better protection against viruses. Furthermore, we found that B-cells constituted the larger human hematopoietic compartment in the thymus of both huNSG-DKO and huNRG. Thymic B-cells have been accounted for mechanisms of tolerance and autoimmunity (Castañeda *et al*, 2021). Although their role is still unclear, since B-cells express high levels of MHC class I/ II and of co-stimulatory ligands, they are well fitted as antigen-presenting cells. We postulate that high number of B-cells could have also contributed to T-cell development in thymus of huNSG-DKO mice. Besides the thymus, we found high amount of T-cells in bone marrow and spleens of huNSG-DKO. Interestingly, in both tissues these T-cells were activated and showed memory phenotypes. Notwithstanding, although T-cells are crucial for T-dependent B-cell maturation (Lang *et al*, 2013), we did not observe substantial differences of mature splenic B-cell compartments of humanized NRG and NSG-DKO mice.

Work by other groups has explored matching the HLA alleles of human cells with transgenic HLAs expressed in mouse tissues. Thus, transgenic mice such as NOD.Cg-*Rag1^tm1Mom^ Il2rg^tm1Wjl^* Tg (HLA-DRA, HLA-DRB1*0401) 39-2Kito/ScasJ (DRAG) and NOD.Cg-*Rag1^tm1Mom^ Il2rg^tm1Wjl^* Tg (HLA-A/H2-D/B2M) DvsTg (HLA-DRA, HLA-DRB1*0401) 39-2Kito/J (DRAGA) were developed. HPCs partially matched to the DRAG/ DRAGA mice (expressing HLA-DR4 and/or HLA-A2.1 genotypes) showed significantly higher reconstitution of T- and B-cells in comparison to non-matched HPC controls (Danner *et al*, 2011; Majji *et al*, 2016; Masse-Ranson *et al*, 2019; Suzuki *et al*, 2012). However, obtaining matched HPCs for a particular transgenic mouse strain is challenging, time consuming and expensive, giving the number of combinations of different HLA haplotypes. Recently, we showed an alternative lentivirus-based approach to deliver the expression of HLA-DR4. LV were used systemically to introduce HLA-DR4 and hGM-CSF/hIFN-α/HCMV-gB in huNRG mice, enhancing B-cell development compared to animals that did not receive the LV injection (Kumar *et al*, 2021). Therefore, a plausible explanation is that the NRG background seems to be fitter to promote B cell development than the NSG strain. In our current study, we also used LVs to introduce HLA-DR4 and HLA-A2.1 into mice humanized with HLA-A2.1^+^ HPCs. Additionally, mice were infused with LV expressing hGM-CSF/hIFN-α/HCMV-gB to boost human adaptive immunity. We observed that detection of human leukocytes was enhanced upon LV administration, which is assumed to be due to HLA matching between CD34^+^ HPCs and *in vivo* transduced mouse cells. One related study compared transgenic NSG mouse strain expressing HLA-A2 with NSG mice injected i.v. or intrathoracically with adeno-associated virus (AAV) expressing HLA-A2 (Huang *et al*, 2014). HLA-A2 expression in thymus was confirmed for both groups by immunohistochemistry and was associated with increased HIS development in comparison to parental NSG mice not expressing HLA-A2. We observed higher T-cell levels in spleen and bone marrow of huNSG-DKO+LV mice in comparison to animals that were not treated with LV. We believe that the superior humanization obtained through LV administration may be also related to a vaccination effect *per se,* leading to an inflammatory milieu promoting higher development of human monocytes and dendritic cells.

Furthermore, using CyTOF, we also detected natural killer cells and γδ T-cells in huNSG-DKO mice, which are minor but relevant participants in immune responses against viruses and tumors. In fact, detection of higher levels of human myeloid and natural killer cells in humanized mice has been associated with superior human immune responses (Douam *et al*, 2018). Further, we observed enhanced expression of CD69 in neutrophils of huNSG-DKO+LV. Since it was shown that GM-CSF can activate neutrophils and enhance CD69 expression (Atzeni *et al*, 2002), this seems to confirm that tricistronic LV promoted an immunization boost.

In line with the immunophenotypic observations, the transcriptomic data showed several correlations with the flow cytometry and CyTOF: higher humanization and activation of immune responses in huNSG-DKO mice. In particular, mRNA expression analyses highlighted the functional activation of several pathways of human adaptive immunity, such as upregulation of genes responsible for defense against viruses and other organisms and higher expression of polyclonal TCRs.

As several human lymphocyte populations require species-specific cytokines to develop and grow, we anticipate that NSG-DKO (and NRG-DKO) mice expressing human cytokines will be soon available. Taking as an example the nowadays popular NSG-SGM3 (NOD.Cg-*Prkdc^scid^ Il2rg^tm1Wjl^* Tg(CMV-IL3,CSF2,KITLG)1Eav/MloySzJ) mice, expressing human stem cell factor (SCF), granulocyte-macrophage colony-stimulating factor (GM-CSF), and interleukin (IL) 3. This strain is known upon injection of CD34^+^ CB cells to develop not only human T-cells but also a broad myeloid compartment, in particular eosinophils and macrophages (Coughlan *et al*, 2016; Janke *et al*, 2021). Moreover, the MISTRG strain _(Balb/c;129S4-*Rag2*_*tm1.1Flv _Csf1_tm1(CSF1)Flv _Csf2/Il3_tm1.1(CSF2,IL3)Flv _Thpo_tm1.1(TPO)Flv _Il2rg_tm1.1Flv* Tg(SIRPA)1Flv/J) expressing IL-3, GM-CSF, and additionally human signal regulatory protein α (SIRPA), thrombopoietin and human homologs of the cytokine macrophage colony-stimulating factor shows high HIS engraftment efficiency and develops human myeloid, natural killer cells and macrophages. Due to heightened replacement of mouse hematopoietic progenitors in the bone marrow and incomplete human erythropoiesis, the mice can commonly become anemic and die (Rongvaux *et al*, 2014). Furthermore, humanized NSG-FLT3 (NOD.Cg-*Flt3^em2Mvw^ Prkdc^scid^ Il2rg^tm1Wjl^*Tg(FLT3LG)7Sz/SzJ) mice with human FLT3 ligand knocked in and mouse FLT3 receptor knocked out have significantly higher levels of human monocytes, dendritic, natural killer and T-cells in comparison to NSG mice (Yao *et al*, 2022). A similar effect was seen also in different strain but with the same transgenic mutations to diminish mouse FLT3 and introduce human FLT3 (Li *et al*, 2016). Thus, NSG-SGM3 and NSG-FLT3 strains showing superior humanization and development of different populations of myeloid cell compartment would be promising candidates to be cross-bred with NSG-DKO mice.

In summary, huNSG-DKO mice showed high humanization and the absence of mouse MHCs did not hinder T-cell development. This contradicted the paradigm that human T-cells have to be educated on mouse epithelial MHCs in humanized mouse models, and opens new perspectives regarding the roles of B-cells or dendritic cells to participate in the positive and negative T-cell selections. Ultimately, as HIS mice are becoming part of several translational programs to test or compare immunotherapies against cancer, the next experimental challenge will be to match the HLAs of the HPCs, HLAs expressed in mouse and HLAs in the tumor. Towards this goal, artifacts caused by xeno-GvHD can be potentially reduced by using the NSG-DKO strain. Further, once the HLA matching is optimized or completed, higher levels of antigen-specific T-cell responses can be measured. All in all, our proof-of-concept study will contribute to the future development of optimized NSG-DKO humanized mice for broad uses in basic immunology and translational research.

## Methods

### Lentivirus production

The self-inactivation third generation lentivirus vectors LV-HLA-DR4/fLuc, LV-HLA-A2.1/fLuc and LV-hGM-CSF/hIFN-α/HCMV-gB (**Figure EV4**) were produced and tittered as previously described (Theobald *et al*, 2020; Kumar *et al*, 2021).

### HPC selection

The study was conducted according to the guidelines of the Declaration of Helsinki, and approved by the Ethics Committee of the Hannover Medical School (MHH). Umbilical CB samples were collected after informed consent from the mothers at the Department of Gynecology and Obstetrics (MHH, Hannover) and obtained according to study protocols approved by MHH Ethics Review Board (approval obtained on 12 December 2012, number 4837 to RS). After CB collection, CD34^+^ cells were positively selected through two consecutive runs using immune magnetic beads kits (Miltenyi Biotec, Bergisch-gladbach, Germany) and cryopreserved, and only units with more than 98% purity (by FACS) were used for studies as previously described (Daenthanasanmak *et al*, 2015). All cryopreserved CB and tissue samples were transferred to the University Hospital of Cologne after approval of the ethics committee.

### Generation of humanized mice

The animal protocols for mouse studies were approved by the Lower Saxony Office for Consumer Protection and Food Safety–LAVES (approval number 33.19-42502-04-19/3336 to RS) and performed according to the German animal welfare act and the EU directive 2010/63. The well-being of mice was monitored according to score sheets approved by LAVES with pre-defined humane endpoints. Breeding pairs of NRG mice (stock number 017914, NOD.Cg-*Rag1^tm1Mom^ IL-2rγc^tm1Wjl/SzJ^*) and NSG-DKO (stock number 025216, *NOD.Cg-Prkdc^scid^ H2-K1^b-tm1Bpe^ H2-Ab1^g7-em1Mvw^ H2-D1^b-tm1Bpe^ IL-2rgc^tm1Wjl/SzJ^*) were obtained from The Jackson Laboratory (JAX; Bar Harbor, ME, USA) and bred and maintained in house under pathogen-free conditions. For some experiments, NSG-DKO mice were directly purchased from The Jackson Laboratory. Briefly, 5–7-week-old mice were sub-lethally irradiated (450 cGy for NRG, 150 cGy for NSG-DKO) using a [^137^Cs] column irradiator (Gammacell 2000 Elan; Best Theratronics, Ottawa, ON, Canada), and 4 h after irradiation, 1–2 x 10^5^ human CB CD34^+^ cells were injected i.v. into the tail vein of mice. CB units were tested prior to experiments for ascertaining their human immune reconstitution. Only CB units that resulted in >20% huCD45^+^ cells in PBL at week 10 post-HCT were used in studies. The animals received antibiotics (Cotrim-K, Ratiopharm) in their water two days prior irradiation and continued receiving it for 14 days post-HCT. The body weights and general health of the mice were monitored three times per week after HCT.

### *In vivo* administration of LVs into mice and BLI analyses

HuNSG-DKO mice were divided into two groups. One of the groups received LV while other did not. In detail, 1 µg of p24 equivalent of each LV were injected i.v. into the tail vein. At week 1 post-HCT HLA-DR4/fLuc, HLA-A2.1/fLuc were administered and week 8 post-HCT hGM-CSF/hIFN-α/HCMV-gB was injected. To visualize fLuc expression, mice were analyzed at week 8 and 12 post-HCT by BLI analyses using the IVIS SpectrumCT (PerkinElmer, Waltham, MA, USA) as described (Theobald *et al*, 2020). Briefly, mice were anesthetized using isoflurane and shaved. Five minutes before imaging, mice were injected i.p. with 2.5 µg D-Luciferin potassium salt (SYNCHEM, Elk Grove Village, IL, USA), freshly reconstituted in 100 µL PBS. Images were acquired in a field of view C, f stop 1, and medium binning for each mouse. Exposure time was kept to 300 s for each mouse. Data were analyzed using LivingImage software (PerkinElmer, Waltham, MA, USA).

### Blood and tissue collection and processing

Immune-reconstitution in the PBL was monitored at 8, 12, 16 and 20 post-HCT. At the endpoint analyses (week 20 post-HCT), PBL was collected and several tissues were biopsied (spleen, bone marrow, and thymus) and processed as previously reported (Theobald *et al*, 2020). The tissues were cryopreserved in cryo-medium (40% PBS; 50% Human Serum, Sigma-Aldrich, St. Louis, MO, USA; and 10% DMSO) and stored at -150 °C for further analysis.

### Blood analysis by flow cytometry

PBL samples were incubated with lysis buffer (0.83% ammonium chloride/20 mM HEPES, pH 7.2) for 5 min at room temperature to remove erythrocytes. Samples were then blocked in PBS plus 10% FBS and stained with optimum concentration of antibodies for flow cytometry (**Suppl. Table 2**), and additional washing was performed to remove unbound antibodies. For data acquisition, an LSR II flow cytometer (BD Biosciences, Heidelberg, Germany) was used and analysis was performed using FlowJo software (Treestar Inc., Ashland, OR, USA). Gating strategies can be seen in **Figure EV2A**. The data was visualized using GraphPad Prism software (Dotmatics, Boston, Massachusetts).

### Tissue analysis by flow cytometry

Cyropreserved single-cell suspensions were blocked with mouse IgG block (Sigma Aldrich, USA) and human FcR binding inhibitor (eBioscience). Dead cells were excluded using Zombie UV dye (Biolegend). Afterwards, cells were stained for flow cytometry for 20 min at 4 °C using surface antibodies (**Suppl. Table 2**). Data were acquired on a Cytoflex LX flow cytometer (Beckman Coulter, Brea, California, USA) and analyzed using the Kaluza software (Beckman Coulter, Brea, California, USA). Gating strategies can be seen in **Figure EV2B, C and EV3A, E**.The data was visualized using GraphPad Prism software (version 9.5.0).

### CyTOF

The bone marrow cells were thawed then stained and barcoded with CD45-Cd using Stardard Biotools protocols (**Suppl. Table 7**). The samples were measured using HELIOS, a CyTOF system (Standard BioTools, South San Francisco, California, USA).

Signal intensity measured in a mass cytometry (CyTOF) channel is often susceptible to interference from neighboring channels due to technological constraints. These interferences, known as spillover effects, can significantly hinder the accuracy of cell clustering. Most of the current approaches mitigate these effects through the use of additional beads for normalization, known as single-stained controls. However, this method can be costly and necessitates a customized panel design. To address these challenges, we employed CytoSpill (Miao *et al*, 2021), a tool that quantifies and compensates for spillover effects in CyTOF data without the need for single-stained controls. Mass cytometry spillover-corrected data were transformed using an inverse hyperbolic sine (arcsine) function (Diggins *et al*, 2015) with a co-factor of 5. We further apply a 99.9% marker normalization step, where each areasinus hyperbolicus (arcsinh)-transformed marker is normalized by its 99.9th percentile value. For cell type identification, we carried out a two-step process. First, we performed unsupervised clustering of cells, followed by assigning cell types to each cluster. After defining a subset of relevant markers, we utilized the FlowSOM (Van Gassen *et al*, 2015) clustering algorithm, incorporating Self-Organizing Map clustering and Minimal Spanning Trees, to cluster all cells into 100 groups based on the expression of lineage-defining markers presented in the dot plots. Subsequently, we metaclustered these initial clusters into 10 biologically relevant clusters using consensus hierarchical clustering. The final clusters underwent manual refinement and annotation based on the median expression profile of individual metaclusters. Clusters containing non-biologically meaningful signals were labeled as ‘Unassigned’. Spillover correction was conducted utilizing the CytoSpill R package accessible on GitHub at https://github.com/KChen-lab/CytoSpill. The clustering algorithm FlowSOM (Bioconductor FlowSOM package in R) is available at https://github.com/SofieVG/FlowSOM. Plots were created in python using scanpy (Wolf *et al*, 2018) version 1.9.3 which is a scalable toolkit for analyzing single-cell data. For data visualization, high-dimensional single-cell data were reduced to two dimensions using the nonlinear dimensionality reduction algorithm t-SNE (Maaten & Hinton, 2008). tSNE plots were created and visualized using scanpy. The barplots were created using GraphPad Prism software (Version 9.5.0).

### mRNA sequencing and bio-informatic analyses

Spleen cells that were not used for FACS analysis were resuspended in 1 mL Trizol (Invitrogen, Waltham, Massachusetts, USA) and frozen at -80 °C until further use. The RNA was extracted using RNeasy MinElute kit (Qiagen, Hilden, Germany). In detail, the cells frozen in Trizol were thawed on ice and 200 µL chloroform added. The samples were centrifuged at 12,000 g for 12 min at 4 °C. Then aqueous phase was transferred to a new tube and the same volume of 70% ethanol was added. The mixture was transferred to RNeasy MinElute spin column and centrifuged at 8,000 g for 15 second at RT. The flow-through was dumped and 500 µL RPE buffer added. The samples were once again centrifuged at 8,000 g for 15 seconds at room temperature and flow-through was dumped. Then 500 µL 80% ethanol was added and the samples were centrifuge at 8,000 g for 2 min at RT. Afterwards, the samples were dried by centrifugation at max speed for 5 min. The RNA was extracted using 14 µL RNase-free water and centrifugation at max speed for 1 min. The quantity and quality of RNA was checked using photometer (NanoPhotometer N60, Implen, Munich, Germany). NGS analyses were carried out at the production site Cologne (Cologne Center for Genomics (CCG)). Alignment was performed in two different approaches. First, we mapped all samples to the mouse (GRCm39) and human (GRCh38) genomes separately using the Nextflow rnaseq pipeline (3.12.0) to evaluate the mouse and human specific samples for QC. Second, as the humanized mice samples represent a DualSeq approach (measuring two separate species simultaneously) we combined the human and mouse genomes and repeated the mapping using the same Nextflow rnaseq pipeline but with some changes to the parameters to the STAR aligner. Specifically, we adopted the ENCODE standard options for the STAR aligner in addition to modifying the “outFilterMultimapNmax” parameter from 20 to 40 as we expected many potential overlapping mappings between the two fairly homologous genomes. All analyses were performed using DESeq2 with the output of the Nextflow rnaseq pipeline. The DESeq models were only evaluated against the two groups compared. Identified differentially expressed genes were then first separated into mouse and human specific genes and then functionally annotated using the gprofiler2 package. Resulting Gene Ontology Biological Processes were then further processed using the rrvgo package to derive semantic similarity plots for further interpretation.

### Statistics

For the blood values, 3-Way-ANOVA with categorical variables mouse, week and treatment group was applied in order to test whether the repeated measures reflected by the variable Mouse had an influence. Since there was no significant influence by the repeated measures, pairwise t-tests were applied with Bonferroni-Holm correction for multiple testing. For organ data (FACS and CyTOF) with more skewed distributions, the Wilcoxon-Mann-Whitney test was applied also with Bonferroni-Holm correction. Statistical analysis was carried out with the statistical software R version 4.2.3. All calculated p-values can be found in **Suppl. Table 4 - 6** for FACS and **Suppl. Table 8, 9** for CyTOF data.

## Data availability

The datasets and computer code produced in this study are available in the following databases:

- **RNA-Seq data:** data are available in the BioStudies database (http://www.ebi.ac.uk/biostudies) under accession number E-MTAB-13883.

## The Paper Explained

**Problem:** The major histocompatibility complexes (MHCs) enable development of adaptive immunity through presentation of peptidic antigens to T-cells. Multi-allelic MHCs are the most polymorphic genetic system in humans. MHCs are immunogenic *per se* and determine the feasibility of transplantations. Despite the full MHC mismatch, hematopoietic human stem cell transplantation into immune deficient mice results into humanized mice with human T, B and myeloid compartments. Humanized mice are explored to test human-specific immune responses against pathogens and cancer. However, the mechanism of cross-talk between the mouse and human MHCs systems and human T-cell development have not been fully elucidated. Here, we tested if human T-cells require the expression of mouse MHCs in tissues for their differentiation and activation.

## Results

Using a novel commercially available immunodeficient mouse strain lacking expression of MHC class I and II, we observed a rapid human hematopoiesis and achieved development of human T-cells in thymus and several lymphatic tissues. We also found that a vaccine delivering human MHCs, cytokines and a herpesvirus antigen boosted the development of polyclonal αβ and γδ T-cells in humanized mice. In addition, vaccination upregulated the expression of genes involved in multiple pathways of human immune reactivity.

### Impact

These findings revealed that this novel mouse strain may improve humanization and that donor T-cell development *in vivo* does not require expression of MHCs in the recipient. Future research exploring replacement of the murine MHC expression in recipient tissues and delivery of matched HLA alleles is warranted to limit the xenograft or allograft reactivities and improve the outcome of inter- and intra-species transplantations.

### For More Information

For more information about humanized mice and EMBO Practical Courses visit: https://www.jax.org/jax-mice-and-services/in-vivo-pharmacology/humanized-mice

Innovations, challenges, and minimal information for standardization of humanized mice | EMBO Molecular Medicine (embopress.org)

EMBO Practical Course Humanized Mice in Biomedical Research - Sunday 15 October-Friday 20 October 2017 (lifescience.net)

Humanized Mice in Biomedicine: Challenges and Innovations – Course and ConferenceOffice (embl.org)

Humanized mice, personalized therapies and big data – Course and ConferenceOffice(embl.org)

Humanized mice: immunotherapy and regenerative medicine – Course and ConferenceOffice (embl.org)

## Acknowledgments

We thank James Keck from The Jackson Laboratory Sacramento for his great support to get this project started. We thank Dirk Wedekind (Animal Facility, Hannover Medical School), who kindly assisted with the preparation and submission of animal study protocols. We thank all the staff of the Clinic for Obstetrics of the Hannover Medical School for procurement of CB. We thank Rainer Blasczyk of the Institute for Transfusion Medicine and Transplant Engineering for providing the HLA genotyping upon recharge. We thank the Cologne Center for Genomics at the University of Cologne for sequencing the mRNA. The authors thank members of the Regenerative Immune Therapies Applied Laboratory at the Hannover Medical School and the Institute of Translational Immuno-Oncology at the University of Cologne for kind technical support.

## In Memoriam

We sincerely thank Dr. Suresh Kumar, who is in our thoughts, for his hard work, kind dedication and highly important contributions to this work.

## Author Contributions

M.D. and J.K.curated the samples and data, completed the flow cytometry analyses. M.D. prepared the samples for CyTOF and mRNA sequencing and prepared the final versions of the figures, wrote the first draft and assisted in the editing. J.K. and A.S. produced and tittered LVs. J.K., T.B. and A.S. performed *in vivo* experiments and collected and cryopreserved tissues. J.K. and T.B. performed flow cytometry measurements of blood. M.D., H.S., K.W., M.T., S.J.T. designed T and B panels and performed flow cytometry measurements of organs. D.B. and P.H. stained and measured samples on CyTOF. N.A. and R.G. performed bioinformatics analysis of CyTOF data. P.A. analyzed RNAseq data. M.R. performed bioluminescence analyses of tissues. J-M.K. assisted with the TCR analyses and deposit of data. F.K. performed the statistical analysis. CvK provided the CB under informed consent of the mothers. S.T. Provided the power calculations for experimental design. M.G.M and E.B. provided the information about HLA typing. M.D. and R.S. administered the resources and financials of the project. R.S., B.S., L.S. performed the conceptualization of the studies, obtained the funding and interpreted the data. R.S. obtained the regulatory approvals, coordinated the interactions of the project and participated in the composition of the first draft and revisions of the final manuscript. All authors have read and agreed to the published version of the manuscript.

## Expanded view figure legends

**Figure EV1:**

A Relative weight of huNRG (black), huNSG-DKO (blue) or NSG-DKO+LV (red) after reconstitution with CD34^+^ isolated cells from three different CB donors. HCT was performed at day 0 and mice were sacrificed at day 120. The weight measured at day 0 was considered the 100% reference for each mouse. For the NSG-DKO+LV cohort (red), LVs were administered on week 1 (day 7) and week 8 (day 56) after HCT.

B Photographs corresponding to detection of bioluminescent signals in non-humanized NSG-DKO mice injected with LV-DR4/fLuc and LV-A2.1/fLuc. The control mouse was injected with PBS. BLI measurements were performed at 8 weeks post LV or PBS injection. C Quantification of BLI signal of humanized mice injected with LV-DR4/fLuc and LV-A2.1/fLuc. The control mouse (black bar) was injected with PBS. BLI measurements were performed at 8 and 12 weeks post-HCT. Quantification shows results for three different mice injected with LV (grey bars).

**Figure EV2:**

A Gating strategy for flow cytometry analyses of human leukocytes in blood samples. Example showing blood of a huNSG-DKO+LV mouse as a reference.

B Gating strategy for flow cytometry analyses CD45^+^ and CD34^+^ cells. Example showing spleen of a huNSG-DKO+LV mouse as a reference.

C Gating strategy for flow cytometry analyses of human CD45^+^ and CD34^+^ cells. Example showing CD34^+^ cells selected from cord blood as reference.

D Flow cytometry analyses of bone marrow showing quantification of cells expressing huCD45, huCD3 or huCD34 (in absolute cell counts, log scale).

E Flow cytometry analyses of thymus and quantification of cells expressing huCD4, huCD8, huCD4/CD8, huCD19 (in absolute cell counts, log scale).

F Flow cytometry analyses of spleen and quantification of cells expressing huCD4, huCD8, huCD4/CD8, huCD19 (in absolute cell counts, log scale).

**Figure EV3:**

A Gating strategy for flow cytometry analyses and detection of PD-1 and CD69 activation markers. Example showing spleen of a huNSG-DKO mouse as reference.

B Analysis of Naive and Terminal Effector T-cell subtypes within huCD4^+^ and huCD8^+^ T-cells in bone marrow (in percentages).

C Analysis of Naive and Terminal Effector T-cell subtypes within huCD4^+^ and huCD8^+^ T-cells in thymus (in percentages).

D Analysis of Naive and Terminal Effector T-cell subtypes within huCD4^+^ and huCD8^+^ T-cells in spleen (in percentages).

E Gating strategy for flow cytometry analyses of B-cell subtypes. Example showing cells recovered from spleen of a huNSG-DKO+LV mouse as reference.

F Enumeration of IgM^+^, IgM^-^IgA^+^ and IgM^-^IgG^+^ B-cells in spleen (in absolute cell counts, log scale).

**Figure EV4:**

A Schematic representation of the multicistronic lentiviral vectors used in the study.

B Detection of HLA-DR4 (upper panel) and HLA-A2.1 (lower panel) on 3T3 mouse fibroblasts transduced with lentiviral vectors.

**Figure EV5:**

A CyTOF analyses to validate the methodology with reference samples: human cord blood (left panel), bone marrow of non-humanized NRG mouse (middle panel) and bone marrow of non-humanized NSG-DKO mouse (right panel).

B CyTOF analyses and dotplot of huNRG and huNSG-DKO cell subtypes and their expression of different Maxpar Direct Immune Profiling lineage markers. The dot size corresponds to the fraction of cells expressing the indicated marker within each cell type, and the color indicates median expression.

